# Widespread genome instability in *Solanum tuberosum* plants regenerated from protoplasts

**DOI:** 10.1101/382861

**Authors:** Michelle Fossi, Luca Comai

## Abstract

Non-transgenic genome editing in regenerable protoplasts, cell-wall free plant cells, could revolutionize crop improvement because it reduces regulatory and technical complexity. But, plant tissue culture is known to engender frequent unwanted variation. To evaluate the contribution of genome instability to this phenomenon, we analyzed large scale copy number changes in potatoes regenerated from protoplasts by comparison of Illumina read depth. While a control set of plants that had been propagated by cuttings displayed no changes, the protoplast regenerants were affected by pervasive aneuploidy. In addition, certain chromosomes displayed segmental deletions and duplications ranging from one to many. Resampling the same plant found different dosage profiles in different leaves, indicating frequent persistence of instability.

## Introduction

Protoplast modification via nucleoprotein complexes results in high efficiency editing and transgene-free genomes (Woo et al., 2015; Andersson et al., 2018). If regeneration is possible, at least some of the resulting plants should display only the targeted changes. By contrast, regulatory compliance when using stably integrated transgenes requires, at a minimum, elimination by meiotic segregation. This, however, is not a viable approach in clonally propagated, highly heterozygous crops because the optimal parental genotype is unlikely to be frequent in the progeny. Somaclonal variation, i.e. genotypic and phenotypic differences from the source plant (Landsmann and Uhrig, 1985; Lee and Phillips, 1988; Bao et al., 1996), has been associated to epigenetic changes (Stroud et al., 2013; Ong-Abdullah et al., 2015; Han et al., 2018), single nucleotide mutations (Miyao et al., 2012), structural changes, and aneuploidy (Lee and Phillips, 1988). Notwithstanding the high likelihood of large phenotypic impact, chromosomal alterations remain poorly documented. Potato *(Solanum tuberosum)* is a major caloric source in both industrialized and developing countries. Its breeding is complicated by its autotetraploid and highly heterozygous genome, which also hinders the use of sexual seed (Potato Genome Sequencing Consortium et al., 2011; Hardigan et al., 2017). In the 80’s, potato emerged as a model for somaclonal variation, even spurring commercial interest as an alternative to breeding (Evans, 1989; Karp et al., 1989). Multiple varieties were shown to be amenable to protoplasting followed by regeneration. Notwithstanding the interest, production of relevant commercial varieties through this method has been limited (Krishna et al., 2016). The possibility of chromosomal instability was highlighted in cytological studies, but structural changes were detected infrequently (Karp et al., 1982; Creissen and Karp, 1985; Sree Ramulu et al., 1986).

Interestingly, the outcome of the regeneration process was thought to result in a binary outcome: either somaclonal variants, or normal plants that resembled in all characteristics the protoplasted variety. Implicitly, the assumption was that the latter plants had dodged the “somaclonal bullet”. Lacking a mechanistic understanding of what causes somaclonal variation, this assumption is often extended to explant regenerants used for Agrobacterium or biolistic transformation. In other words, off-types commonly encountered among plants produced by dedifferentiation and proliferation of somatic cells may be the product of mutagenic mechanisms whose frequency can be minimized and phenotypic screens are adequate to address the problem. The need to explore these assumptions is underscored by the unexplained yield drag connected with transgenic modifications (Elmore et al., 2001). To enhance our molecular understanding of somaclonal variation, we studied copy-number changes in a set of regenerated potatoes demonstrating an unexpected degree of chromosomal and segmental instability in their genomes.

## Results and Discussion

We produced protoplasts from autotetraploid potato variety Desiree and induced regeneration through callus and shoot formation (Fig. 1-A). In preparation for genome editing, we wanted to measure the rate of sterility and abnormalities among regenerated plants. In two experiments we collected ~ 400 plants, some from the same callus and thus protoplast. Of 101 plants transplanted to the greenhouse most tuberized bypassing flowering (Suppl. Table 1 and Suppl. Table 2). Of the 26 that flowered, 25 had viable pollen (Suppl. Table 3). Five displayed abnormal morphology (Suppl. Table 4, Suppl. Fig. 1). To investigate whether chromosomal alterations were associated with these abnormalities, we analyzed the genomic DNA of each individual. We also selected 10 additional regenerated plants that appeared normal.

**Figure 1.**
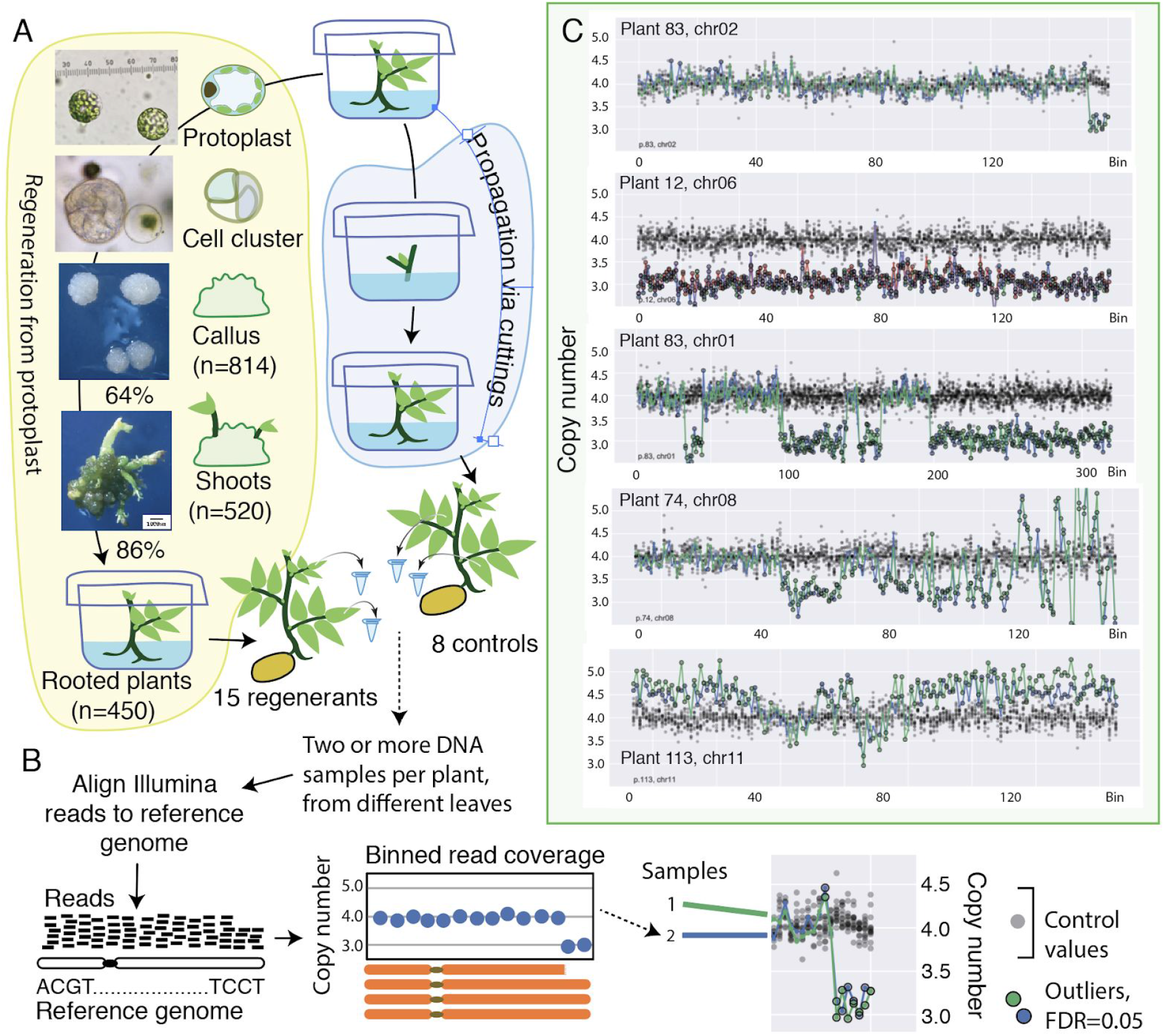
Plant production and analysis. A. Autotetraploid potato var. Desiree was cultured axenically and either protoplasted and regenerated (left) or propagated from nodal buds without callus formation or regeneration (right). Cumulative numbers or efficiencies for two experiments are shown. B. Derivation of chromosomal dosage and plot display. C. Dosage plots illustrating variation for selected chromosomes. Two or three sample values from a single regenerated plant are plotted over the controls (black dots). Four genomic copies are expected from autotetraploidy.

Illumina sequence reads from genomic DNA were mapped to the potato reference genome (Hardigan et al., 2016). Read counts (mean=1155; std=268) binned in 0.25Mb consecutive but non-overlapping genomic bins were normalized for chromosome copy dosage using the counts from a single propagated control plant (Fig. 1-B; see Methods on line). *S. tuberosum* has 12 chromosomes, each in four copy (tetrasomy). Eight control plants propagated by nodal cutting without protoplasting or regeneration, and their replicate samplings, displayed regular genomes (Suppl. Fig. 2).

Standardized read coverage of regenerated plants was compared to controls, identifying outliers. In each plant, multiple bin measurements differed from the expectation of four copies. They encompassed segments ranging from a few bins to whole chromosomes (Fig. 1-C). Reliability of each structural variant identification increases with the number of contiguous outliers enabling robust conclusions.

Cloning by culturing stem cuttings on artificial media, a common horticultural practice, did not cause genome instability. The process of protoplast regeneration, however, engendered high instability, which affected both normal and abnormal regenerated plant types. Among the 15 regenerated plants (Fig. 2-A) those phenotypically abnormal had more changes affecting whole chromosomes, (2.83 vs 0.9; P = 0.0035, 2-tailed T-test). However, mean occurrence of any change (small or large) did not differ between normal appearing and abnormal cohorts, suggesting that screening for normal individuals may not select for plants with intact genomes.

**Figure 2.**
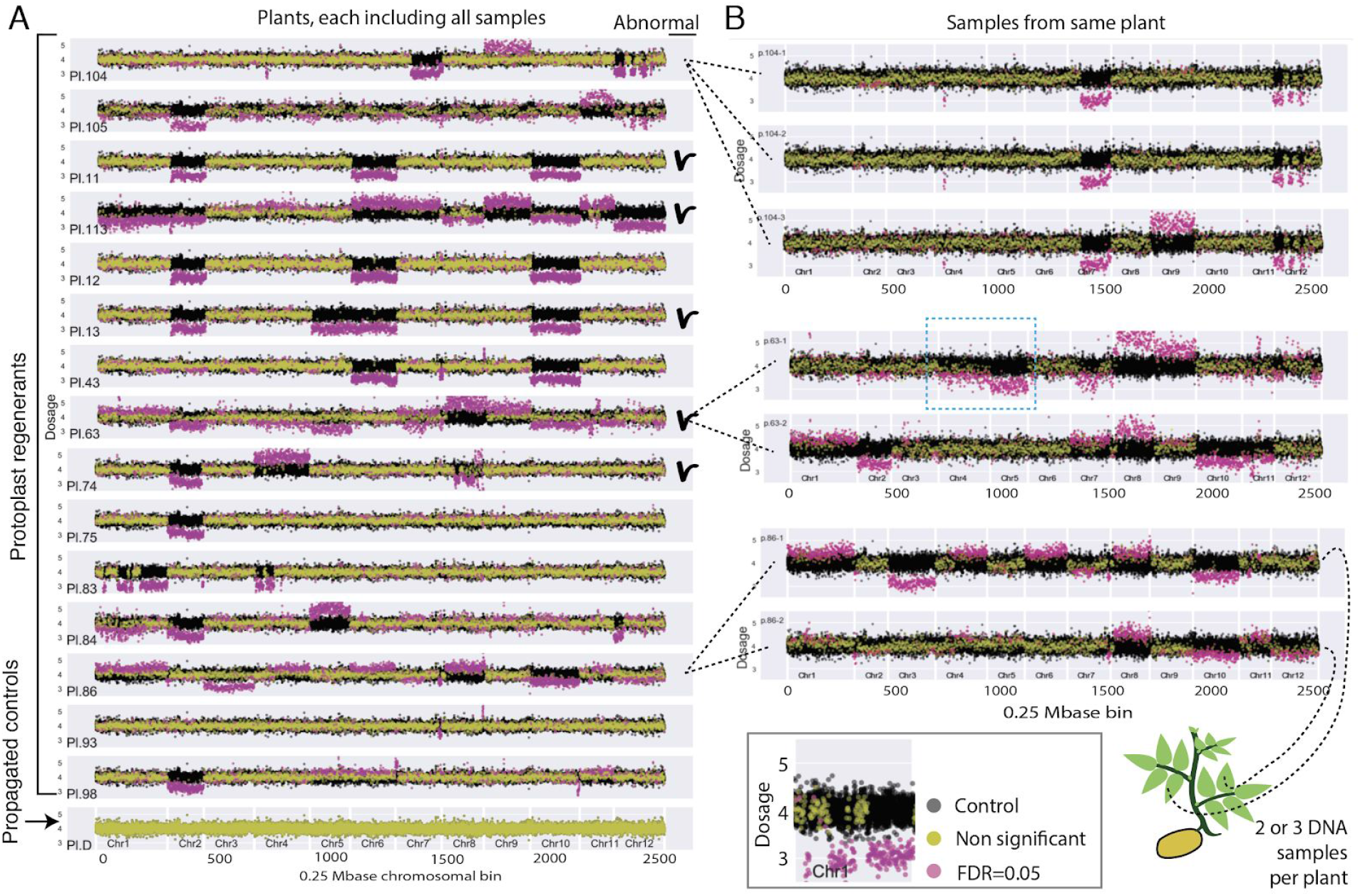
Genome dosage changes in plants regenerated from protoplasts. Values on y-axis are plotted versus 0.25Mb chromosomal bins on the x-axis, arrayed consecutively for the 12 chromosomes of potato. Control data are in black. Four genomic copies are expected from autotetraploidy. Bins with high variability were dropped (see Methods). A. Two to four independent samples are plotted together for each plant, except for P.105. Since calli could be resampled, some plants may derive from the same protoplast. B. Individual plotting of discordant replicate samples from different leaves of the same plant, indicating persistent instability. Some chromosomes display incomplete shifts (blue hatched box, additional cases visible in A and B) consistent with a mix of different karyotypes in the sampled DNA.

After scoring the large multiple-bin changes we concluded that all chromosomes could be affected, but with varying frequency (7-73%). In several instances, a single chromosome (1, 4, 8, and 12) displayed multiple deletions or duplications (Fig. 1-C, Fig. 2-A). This degree of fragmentation resembles chromothripsis, as described in human cancer (Leibowitz et al., 2015) and in plants (Tan et al., 2015). The extent of rearrangements cannot be inferred accurately from our data: at least some of the apparent chromosomal discontinuities could be accounted for if the genome of *S. tuberosum* var. Desiree differs from that of the DM1-3 reference used here for sequence alignment (Suppl. Fig. 3), as demonstrated for the pericentromeric region of chr. 5 (de Boer et al., 2015). Even if multiple deletions have occurred, our analysis does not reveal if the shattered chromosome fragments form a single reshuffled, but syntenic chromosome as in classical chromothripsis (Leibowitz et al., 2015; Tan et al., 2015), or a diaspora of translocations to different chromosomes. In any case, our results indicate that genome instability results both in aneuploidy and in dsDNA breaks, likely leading to rearrangements upon repair.

Genome instability may be catastrophic or chronic. Three plants displayed variations between samples originating from different leaves of the same plant (Fig. 2-B, Suppl. Fig. 2), such as pentasomy 9 in 1 of 3 samples of plant 104. Consistent with the karyotypic chimerism displayed by these plants, incomplete dosage shifts were found in several plants (p63, p84, p86, p98, p113, Fig. 2-A,B, Suppl. Fig. 2). Uniquely among the 96 plants tested for DNA content by flow cytometry, plant 86 displayed peaks consistent with both 4X and 8X DNA content. Chimerism for ploidy type may thus explain the smaller dosage shift for that particular plant. However, none of the other plants exhibited DNA content exceeding tetraploidy. Thus, the intermediate dosage shifts might originate from chimerism in the sampled tissue, which for example could contain a mixture of cells trisomic or tetrasomic for a given chromosome. We concluded that instability could persist through development, generating new variation.

Taken together, our analysis provides compelling proof of large scale, frequent genomic instability as a consequence of regeneration from protoplast in potato. The complete penetrance of this instability syndrome is probably favored in tetraploid potato due to polyploidy buffering (Comai, 2005). Notably, the high frequency of polyploids among regenerants of diploid potato (Sree Ramulu et al., 1986) could result from selection for lower imbalance. In diploid species, monosomy is highly deleterious and trisomy can be fairly deleterious (Henry et al., 2010). Most highly rearranged chromosomes, such as those produced by chromothripsis are similarly deleterious (Tan et al., 2015). We expect that a corresponding analysis in a diploid would find fewer large-scale abnormalities because gross dosage and structural variants will be selected against during regeneration.

Our data provides sequence-based evidence of multiple and frequent changes. What is the mechanistic basis of this instability? In other studies, somaclonal variation has been observed irrespective of whether the samples originated from protoplasts or not (Lee and Phillips, 1988). This and our observation of persistent instability (Fig. 2-B) suggest that protoplasting is unlikely to be the unique cause. The observed syndrome is consistent with failure of one or more of the major mechanisms contributing to genome stability (Lee and Phillips, 1988; Lee et al., 2016; Lee and Seo, 2018). Frequent aneuploidy could be triggered by mitotic malfunction, such as a defective spindle, resulting in missegregation, chromosome loss and perhaps rescue, possibly through restitution of micronuclei. Collapse of genome maintenance in the defective micronuclear environment is thought to result in chromosome-specific shattering (Crasta et al., 2012; Zhang et al., 2015; Ly et al., 2017). The frequent and often stereotypical chromosomal breaks could result from epigenetic upheaval and transposon activation, which have been well documented (Hirochika et al., 1996; Stroud et al., 2013; Ong-Abdullah et al., 2015; Han et al., 2018). Whatever the causes, confirmation of this syndrome in other species will affect the prospects for protoplast utilization. In vegetatively propagated crops, the load of genomic changes is likely to affect the plant phenotype and alter, most likely in a negative way, agronomic performance. In sexually propagated crops, recombination during backcrossing to an agronomically fit parent should offset the negative effect of most genomic changes, except of those tightly linked to any locus of interest. Last, if genomic instability is pervasive and frequent during commonly used procedures that involve dedifferentiation of specialized cells and regeneration, measures to understand its causes and ameliorate the consequences should be undertaken.

## Materials and methods

### Plant regeneration and growth

Cuttings of *Solanum tuberosum* group *Tuberosum* cv. Desiree (hereafter referred to as Desiree) were aseptically maintained in vitro at 16 hr light/ 8 hr dark, 25°C and 40 μmol m-2s-1 light intensity. Cuttings were propagated in magenta cubes on medium containing MS salts (Murashige and Skoog, 1962) and Gamborg vitamins (Gamborg et al., 1968), supplemented with 1.5 mg ml-1 (6μM) silver thiosulfate (STS) (Perl et al., 1988), 3% (w/v) sucrose, 590 mg/l MES, and 0.7% (w/v) agar, pH 5.8. Potato protoplasts were prepared from 1g of leaves pulled from in vitro grown plants following procedures of Tan, with modifications (Tan et al., 1987). Protoplasts were resuspended at 0.4-0.6×10^6^ protoplasts/ml and embedded in an alginate solution as in (Perales and Schieder, 1993), with some medium modifications, including substitution of liquid MS basal medium with liquid 8P medium (Kao and Michayluk, 1975). Callus and shoot regeneration was achieved as described (Shahin, 1984). After 6-10 days, multicellular structures were transferred to callus stimulating medium, TM-3. Microcalli 1-2 mm in size were transferred to TM-4 medium, 16 hr light/ 8 hr dark, 25°C and 40 μmol m^-2^s^-1^ light intensity for shoot regeneration. One to three shoots per callus clump were excised and rooted on propagation medium. This experiment was originally meant to yield a rate for sterility and strongly abnormal phenotypes. Therefore, the callus of origin was not recorded and it is possible that plants with close identification numbers may have originated from the same callus. In two experiments, we regenerated about 400 plantlets, 101 of which were transplanted to the greenhouse. From these, we selected 10 regenerated plants that appeared normal and 5 plants that looked stunted or otherwise abnormal. All plants originated from a different shoot, i.e. correspond to an independent regeneration event. However, since the callus clump of origin was not annotated, they could in theory originate from the same protoplast. At the same time, through axenic nodal cuttings propagation in tissue culture, we produced 8 rooted plants. These were transferred to the same greenhouse to be employed as controls that had not experienced regeneration. All sampled plants were transferred to the greenhouse at the same time and grown in the same environment. Plants were acclimated to greenhouse conditions (16 hr light/ 8hr dark) in flats under a plastic dome for 1 week before transplanting to 1-gallon pots. Plants were allowed to flower and tuber, however greenhouse conditions favored tuber formation over flowering.

### Regenerated plant ploidy detection by flow cytometry

The youngest fully expanded leaf was selected from each greenhouse-grown plant for ploidy assessment by flow cytometry. Leaves were sliced with a sharp razor blade and nuclei were stained with DAPI (4’,6-Diamidino-2-Phenylindole), processed with Sysmex’s (formerly Partec) CyFlow^®^ Space ploidy analyzer and analyzed with CyPad^®^ software.

### Regenerated plant pollen viability testing

Flowers collected for viability testing were stained with fluorescein diacetate (FDA). Working solution was prepared freshly for each experiment from 2 stock solutions combining 5ml of solution 1 (100mg/L Boric acid, 700mg/L CaCl_2_2H_2_O and 200g/L sucrose) with 2-4 drops of solution 2 (2mg FDA/mL acetone), until working solution becomes just slightly cloudy. These were kept on ice. Anthers were submerged in a drop of the FDA stain solution, incubated for 30 minutes in the dark and then examined with a fluorescent microscope.

### DNA extraction

DNA was extracted from 23 plants: 8 controls and 15 regenerated individuals. Young fully expanded leaves of these plants were selected for DNA extraction. At minimum, 2 independent samples were collected from each selected plant. Two 4mm hole punches were taken from leaf tissue and grinded in 500μl buffer with a tungsten bead at 20.0 frequency 1/s for 1 minute in the QIAGEN TissueLyserII®. The plate was rotated and then grinded for an additional 1 minute. Two methods were then used to extract DNA from the 2 independent samples collected from each plant. In one method, DNA was extracted using a QIAGEN DNeasy^®^ plant mini kit, following the manufacturer’s instruction and, in the other method, DNA was extracted using a CTAB DNA extraction protocol (Henry et al., 2015). The two DNA extraction techniques yielded comparable quantity and quality DNAs.

### DNA quantification and library preparation

Concentration of all extracted DNA samples was measured using SYBR Green and processed on an automated plate reader (Tsai et al., 2011). Samples were adjusted to 20ng/μl before 50μl of each DNA sample was sheared by sonication on a Covaris apparatus. The Kappa Bioscience (www.kappabio.com) Hyper Prep Kit was used for library construction following manufacturer methods. Libraries used indexed adapters. They were pooled and sequenced at the QB3 core facility (http://qb3.berkeley.edu/gsl/) at the University of California, Berkeley.

### Sequenced DNA dosage analysis

Sequenced reads were demultiplexed to provide data by plant DNA sample. Reads were trimmed to remove adapter sequences and low-quality data. The trimmed reads were mapped to the DM1-3 v4.04 reference genome (Hardigan et al., 2016) using BWA-MEM (Li, 2014) to determine coverage by chromosome region. Mapped sequence reads were grouped into bins, fixed length chromosomal intervals whose read count was then mapped as 1 point on a sequence read coverage plot. The Bin-by-SAM script was used which created a table of read values by chromosome in a specified magnitude of bins (Henry et al., 2015). Bin sizes of 1M, 0.5M and 0.25M base pairs were examined and 0.25M displayed the data most clearly. Five samples which libraries had yielded low read counts (<10 million reads) were excluded from analysis. The remaining libraries had between ~3 and 13 million reads. Binned read values were filtered to eliminate noisy bins. Preliminary evaluations of raw read coverage in control lines indicated that these were likely euploids and that all eight displayed very similar read distribution by chromosome. One of the propagated control was chosen arbitrarily as dosage reference representing 4 copies of each chromosome. Each bin was standardized to this reference, yielding sequence read dosage between 1 and 8, with 4 being the expected value for a tetrasomic chromosome. Noisy bin were filtered. These values were analyzed using JMP (SAS) and the Python libraries Pandas (McKinney and Others, 2010), Matplotlib (Hunter, 2007), and Statsmodels (Seabold and Perktold, 2010). Chromosome counts were inferred by comparing dosage plots. In a trisomic individual, the dosage of the affected chromosome would appear around 3; in a pentasomic individual, around 5.

Although biological replicates were available for most of the protoplasted samples, we decided to treat these independently once we realized that genome instability persisted in regenerated plants and some biological replicates were different because of the underlying genetics and not because of sampling errors. Instead, for each bin we compared the distribution of the 13 control values to the dosage value of each single sample from the regenerated plants. A Z score was calculated for each value (deviation from mean/std dev). The two-tailed P value of the Z statistics was derived and corrected for multiple tests using the statsmodels’ multipletests ‘fdr_tsbh’ method, yielding Benjamini-Hochberg corrected P and the connected False Discovery Rate with alpha = 0.05 (Benjamini and Hochberg, 1995; Seabold and Perktold, 2010). Statistically indexed data were plotted and compared. Sequence reads have been deposited in the NCBI SRA and are available under accession number ……..(in process).

## Author contributions

M.F. and L.C. conceived the original research plans, designed the experiments and analyzed the data; L.C. supervised the experiments; M.F. performed most of the experiments; M. F. and L. C. completed the writing. L.C. agrees to serve as the author responsible for contact and ensures communication.

## Acknowledgements

We would like to acknowledge Kirk Amundson, Benny Ordonez, Meric Lieberman for help and assistance during this work, and Isabelle M. Henry for editing suggestions. Research was supported by NSF-Plant Genome IOS Grant 144612: Rapid and Targeted Introgression of Traits via Genome Elimination. M.F. was supported by a fellowship from the H.M. Clause Company.

